# Comparative Genomics and Full-Length TprK Profiling of *Treponema pallidum* subsp. *pallidum* Reinfection

**DOI:** 10.1101/841395

**Authors:** Amin Addetia, Lauren C. Tantalo, Michelle J. Lin, Hong Xie, Meei-Li Huang, Christina M. Marra, Alexander L. Greninger

## Abstract

Developing a vaccine against *Treponema pallidum* subspecies *pallidum*, the causative agent of syphilis, remains a public health priority. Syphilis vaccine design efforts have been complicated by lack of an in vitro *T. pallidum* culture system, prolific antigenic variation in outer membrane protein TprK, and lack of functional annotation for nearly half of the genes. Understanding the genetic basis of *T. pallidum* reinfection can provide insights into variation among strains that escape cross-protective immunity. Here, we present comparative genomic sequencing and deep, full-length *tprK* profiling of two *T. pallidum* isolates from blood from the same patient that were collected six years apart. Notably, this patient was diagnosed with syphilis four times, with two of these episodes meeting the definition of neurosyphilis, during this interval. Outside of the highly variable *tprK* gene, we identified 14 coding changes in 13 genes. Nine of these genes putatively localized to the periplasmic or outer membrane spaces, consistent with a potential role in serological immunoevasion. Using a newly developed full-length *tprK* deep sequencing protocol, we profiled the diversity of this gene that far outpaces the rest of the genome. Intriguingly, we found that the reinfecting isolate demonstrated less diversity across each *tprK* variable region compared to the isolate from the first infection. Notably, the two isolates did not share any full-length TprK sequences. Our results are consistent with an immunodominant-evasion model in which the diversity of TprK explains the ability of *T. pallidum* to successfully reinfect individuals, even when they have been infected with the organism multiple times.

**Author Summary:** The causative agent of syphilis, *Treponema pallidum* subspecies *pallidum*, is capable of repeat infections in people, suggesting that the human immune response does not develop sufficiently broad or long-lasting immunity to cover treponemal diversity. Here, we examined the genomes from two blood-derived isolates of *T. pallidum* derived 6 years apart from a patient who had syphilis four times during the same period to understand the genetic basis of reinfection. We found a paucity of coding changes across the genome outside of the highly variable *tprK* gene. Using deep profiling of the full-length *tprK* gene, we found surprisingly that the two isolates did not share any full-length TprK sequences.

## Introduction

*Treponema pallidum* subspecies *pallidum* (hereafter *T. pallidum*) is the causative agent of the sexually transmitted disease syphilis. Rates of primary and secondary syphilis have steadily increased in the past decade in the United States, Europe, Australia, and China [1–3]. With this increasing burden of syphilis, the development of an effective vaccine against *T. pallidum* has become a public health priority [4,5].

Groups disproportionately affected by the re-emergence of syphilis, including men who have sex with men and persons with HIV, are at high risk for repeated infection [6,7]. Reinfection with *T. pallidum* was noted in 13.6% of individuals attending a sexually transmitted infection clinic in Brazil over an 8-year period [8]. In California surveillance data from 2002-2006 in men who have sex with men, 5.9% of men had repeat infection within two years of the initial infection [7]. This ability to reinfect suggests that *T. pallidum* has mechanisms to evade adaptive immunity developed during prior infections, or that immunity rapidly wanes after infection.

Immune evasion of *T. pallidum* in the rabbit model is associated with variability in the 12 genes of the *T. pallidum* repeat gene family (*tpr*) [9–11]. Many of the *tpr* genes encode major surface proteins and are likely targets of the immune system [9,10]. One of the proteins encoded by these genes, TprK, shows remarkable sequence diversity between and within isolates [12,13]. This diversity is found in 7 discrete variable regions, V1-V7 [12,14,15]. Interestingly, in the rabbit model, variation in V1-V7 accumulates over the course of infection and results in the infecting isolate carrying multiple heterogeneous copies of TprK [15–18]. TprK-directed antibodies in rabbits target the variable regions and show minimal cross-reactivity between strains [19]. The development of heterogeneous copies of TprK is likely a mechanism through which *T. pallidum* evades the host’s immune response.

To date, studies using whole genome sequencing (WGS) to examine *T. pallidum* have focused on describing the genomic diversity among isolates from different patients [13,20,21]. These have confirmed that *tprK* shows inter- and intra-isolate variation [13,20]. The *T. pallidum* organisms circulating in a population appear to be closely related and limited variation between them tends to accumulate in specific genomic loci such as *tprK* [13]. It is unclear, however, whether genomic variability is concentrated in these same regions in reinfecting strains. Nor have any studies performed full-length sequencing of *tprK* to link variable regions. Here we deeply profiled genomic changes between two *T. pallidum* isolates that caused repeat infection in the same person.

## Methods

### Source of samples

Blood was collected in 2003 and 2009 from a participant who was enrolled in a study of cerebrospinal fluid abnormalities in syphilis conducted in Seattle, WA. He was diagnosed and treated for secondary syphilis with neurosyphilis on the initial date, and for early latent syphilis on the second date. The study protocol was reviewed and approved by the University of Washington institutional review board, and written informed consent was obtained from all study participants.

### Treponema pallidum isolation and propagation in New Zealand white rabbits

Procedures for animal care and research were approved by the University of Washington Institutional Animal Care and Use Committee in accordance with the Public Health Service Assurance issued by the Office of Laboratory Animal Welfare. For the first isolate (148B), 3mL of fresh blood in a purple-top (EDTA) tube was inoculated into the testes of a male New Zealand White (NZW) rabbit. The rabbit was monitored by serology (FTA-ABS and VDRL) until seroconversion and then the strain was passaged again through a second rabbit to amplify the treponemal load. The duration of the first passage was 92 days, and the second was 39 days. Treponemes were extracted from testes in a mixture of 50% normal rabbit serum (NRS) and 50% saline, then mixed with an equal volume of sterile glycerol, snap-frozen in a dry ice/ethanol bath, and stored long-term in liquid nitrogen. The second strain (148B2) was not isolated immediately. Fresh EDTA blood was mixed with an equal volume of NRS and sterile glycerol, snap frozen, and stored in liquid nitrogen as above. Five years later, 3mL of this mixture was thawed and inoculated into a male NZW rabbit and followed by serology as above, followed by a second passage, which was frozen and stored as above. The duration of the first passage was 67 days, and the second was 26 days.

### Quantification of Treponema pallidum in patient blood

0.5 ml of patient blood in EDTA was mixed with 0.5mL of 2X lysis buffer (20mM Tris-HCl, 0.2M EDTA, 1% SDS) and stored at −80°C until testing. The sample was thawed and extracted with the QIAamp DNA Blood Midi Kit (Qiagen, Valencia, CA) according to manufacturer’s instructions. DNA was precipitated with 2.5 volumes of 100% ethanol, 0.1 volumes of 3M sodium acetate, and 1μL of glycogen overnight at −20°C. DNA pellets were washed twice with 75% ethanol and resuspended in 60μL of molecular grade water.

A portion of the *tp0574* gene was amplified using two different sets of qPCR conditions. 5μL of DNA from the first isolate (UW148B) was amplified in a 20μL reaction with 1X Roche LightCycler TaqMan Master Mix (Roche Life Science, Pleasanton, CA), 0.3μM each of sense (CAA GTA CGA GGG GAA CAT CG) and antisense (TGA TCG CTG ACA AGC TTA GG) primers, and 0.1μM of Universal ProbeLibrary Probe #89 (Millipore Sigma, St. Louis, MO). Amplification was performed in LightCycler capillaries (Roche Life Science) on a LightCycler instrument (Roche Life Science) with the following conditions: 95°C for 10 minutes, followed by 45 cycles of 95°C for 10 seconds, 64°C for 30 seconds, and 72°C for 1 second, with data acquisition during the 72°C step. Results were analyzed using the LightCycler software.

For the second isolate (UW148B2), the same portion of T47 was amplified in a 20μL reaction with 1X TaqMan Fast Advanced Master Mix, 0.3μM each of sense (CAA GTA CGA GGG GAA CAT CG) and antisense (TGA TCG CTG ACA AGC TTA GG) primers, and 0.1μM of TaqMan probe (6-FAM-CGG AGA CTC TGA TGG ATG CTG CAG TT- TAMRA). Amplification was performed in a MicroAmp Optical 384-Well Reaction Plate (Applied Biosystems, Grand Island, NY) on the ViiA 7 Real-Time PCR System (Applied Biosystems, Grand Island, NY) with the following conditions: 50°C for 2 minutes, 95°C for 30 seconds, followed by 45 cycles of 95°C for 5 seconds and 60°C for 20 seconds, with data acquisition during the 60°C step. Results were analyzed using the ViiA 7 software program, and for both isolates the number of copies of *tp0574* per μL was used to calculate the number of *T. pallidum* bacteria per mL of blood.

### Capture sequencing for recovery of whole genomes

DNA was extracted from 200 μL of treponemes suspended in NRS-saline using QiaAmp DNA Blood Mini kit and eluted into 100 μL AE buffer. *T. pallidum* DNA was quantified by real-time Taqman PCR. Each 30 μL of PCR reaction contained 500 nM of primers (F: CAA GTA CGA GGG GAA CAT CGA T; R: TGA TCG CTG ACA AGC TTA GG) and 330 nM of probe (FAM-CGG AGA CTC TGA TGG ATG CTG CAG TT-NFQMGB), 14.33 μL of 2x QuantiTect multiplex PCR mix, 0.65 μL of 2x QuantiTect multiplex PCR mix with ROX, 0.03 unit of UNG and 2μL of DNA. Rabbit CFTR gene was used as a housekeeping gene (F: GCG ATC TGT GAG TCG AGT CTT; R: GGC CAG ACG TCA TCT TTC TT; probe: FAM-CCA AAT CCA TCA AAC CAT CC-NFQMGB). EXO internal control was spiked into the PCR reaction to monitor the PCR inhibition. The thermocycling conditions are as follows: 50°C for 2 minutes, 95°C for 15 minutes and followed by 45 cycles of 94°C for 1 minute and 60°C for 1 minute.

Truseq KAPA HyperPlus kit was used for DNA fragmentation and library construction as well as pre-capture shotgun sequencing of these strains. A custom biotinylated capture probe set (myBaits; Arbor Bioscience) designed against three *T. pallidum* reference genomes (NC_021508, NC_018722 and NC_016848) was applied to enrich library target following the manufacturer’s protocol. After enrichment, the library was purified with 0.8X volumes of AMPure XP beads (Beckman Coulter) and sequenced on a 2 × 300 bp Illumina MiSeq run.

Both isolates were sequenced to an average depth of at least 196X. Paired end reads were trimmed with Trimmomatic v0.38, *de novo* assembled with SPAdes v3.13.0 and visualized with Geneious v11.1.4 [22–24]. We compared the two strains by mapping reads from the reinfecting strain to the completed genome of the initial strain. A minimum depth of 10x and allele frequency of 70% was used to screen for variants. All variants were manually confirmed and each insertion or deletion in a homopolymer was verified by examining reads from both shotgun and capture library sequencing.

### Sequencing of repetitive regions to construct complete genomes

As repetitive regions of the genome were unfortunately excluded from the custom capture probe set, we were unable to produce complete genomes through *de novo* assembly of the sequencing reads. We conducted PCR across the gapped regions between the *de novo* assembled contigs with the primers described in S1 Table using the high-fidelity PrimeSTAR GXL DNA polymerase (Takara). The following PCR conditions were used: 98°C for 2 minutes followed by 35 cycles of 98°C for 10 seconds, 54-59°C for 15 seconds and 68°C for 3-7 minutes with a final elongation at 68°C for 10 minutes.

A full length PCR product could not be obtained for the 7.4 kb region containing *tprE, tprF*, and *tprG*. Four overlapping PCR products were produced by conducting nested PCR with the primers described in S1 Table. First, the entire 7.4 kb region was amplified with the following conditions: 98°C for 2 minutes followed by 10 cycles of 98°C for 10 seconds, 55°C for 15 seconds and 68°C for 7 minutes with a final elongation at 68°C for 10 minutes. The resulting product was purified with the DNA Clean and Concentrator kit (Zymo). Using this product as a template, the four overlapping PCR products were obtained with the following conditions: 98°C for 2 minutes followed by 25 cycles of 98°C for 10 seconds, 53-56°C for 15 seconds and 68°C for 3 minutes with a final elongation at 68°C for 10 minutes.

Libraries for each of the *tpr* PCR products were constructed with the Nextera XT kit (Illumina) and sequenced on either 1×160 bp or 2×300 bp Illumina MiSeq runs. Sequencing reads were processed and analyzed as described above.

PCR products spanning the repetitive regions of *arp* were obtained through a nested PCR using the primers listed in S1 Table. The following conditions were used to amplify the entire *arp* gene: 98°C for 2 minutes followed by 10 cycles of 98°C for 10 seconds, 55°C for 15 seconds and 68°C for 1 minute with a final elongation at 72°C for 5 minutes. The resulting product was purified with the Zymo DNA Clean and Concentrator kit. With this product as a template, the following conditions were used to obtain a PCR product spanning the repetitive regions of *arp*: 98°C for 2 minutes followed by 25 cycles of 98°C for 10 seconds, 55°C for 15 seconds and 68°C for 1 minute with a final elongation at 72°C for 5 minutes. The resulting product was purified from a 1% agarose gel using the NucleoSpin Gel and PCR Clean-up Kit (Takara) and Sanger sequenced.

### Long- and short-read sequencing of tprK

To analyze the heterogeneity of *tprK* in of each of the strains, we separately amplified and sequenced *tprK* using long- and short-read approaches. PCR was conducted with *tprK*-specific primers (S1 Table) appended to 16bp PacBio barcodes using CloneAmp HiFi polymerase mix (Takara) and the following conditions: 98°C for 2 minutes followed by 35 cycles of 98°C for 10 seconds, 62°C for 15 seconds and 72°C for 2 minutes with a final elongation at 72°C for 5 minutes. The resulting 1.6 kb *tprK* amplicon was purified using 0.6x volumes of AMPure XP beads.

For long-read sequencing, library construction and sequencing on a Sequel I SMRT Cell 1M with a 10-hour movie were completed by the University of Washington PacBio Sequencing Services. A total of 77,391 and 8,610 PacBio CCS Q20 reads were recovered for *tprK* amplicons for 148B and 148B2. Short-read libraries were prepared with the Nextera XT kit (Illumina) and sequenced on a 1 × 190 bp Illumina MiSeq run. A total of 176,840 and 192,304 Illumina reads were recovered for *tprK* amplicons for 148B and 148B2.

*tprK* sequencing reads were analyzed using custom Python/R scripts which are available on GitHub (https://github.com/greninger-lab/tprk). PacBio reads between 1400-1800 bp were trimmed of PCR primers using the dada2 preprocessing pipeline and denoised using RAD [25,26]. Illumina reads were trimmed using trimmomatic as described above. *tprK* variable regions from the denoised PacBio and trimmed Illumina reads were extracted by regular expression matching of the flanking constant sequences. These variable regions were translated and protein sequences were counted according to read depth. Variable region sequences with at least 5 reads of support for either Illumina or denoised PacBio reads were analyzed and visualized using ggplot2 [27]. Full-length denoised PacBio reads that contained an intact open reading frame for tprK were included in phylogenetic analyses. Phylogenetic trees on full-length tprK protein sequences were generated using FastTree and visualized using ggtree [28,29]. Pielou’s evenness and Shannon diversity measures for each variable region were calculated using the R package VEGAN [30].

### Data availability

The NCBI accession numbers for the complete genomes of UW-148B and UW-148B2 are CP045005.1 and CP045004.1. The sequencing reads used to generate these genomes, as well as those from short- and long-read sequencing of *tprK*, are available under NCBI BioProject number PRJNA534087.

## Results

### Case and treponemal strains

A 42-year old man with HIV infection presented with secondary syphilis in 2003 (Fig 1). Serum RPR titer was 1:256, CD4 count was 748 cells/μL and plasma HIV RNA measured less than 50 copies/mL. Because his cerebrospinal fluid VDRL was reactive, he was treated for neurosyphilis. Quantitative treponemal load testing of his blood yielded a median of 822 (range: 702 – 1,006) copies of *T. pallidum* DNA per mL of blood, of which 3mL were injected into a rabbit and passed a second time to derive the strain UW-148B.

**Fig 1.**
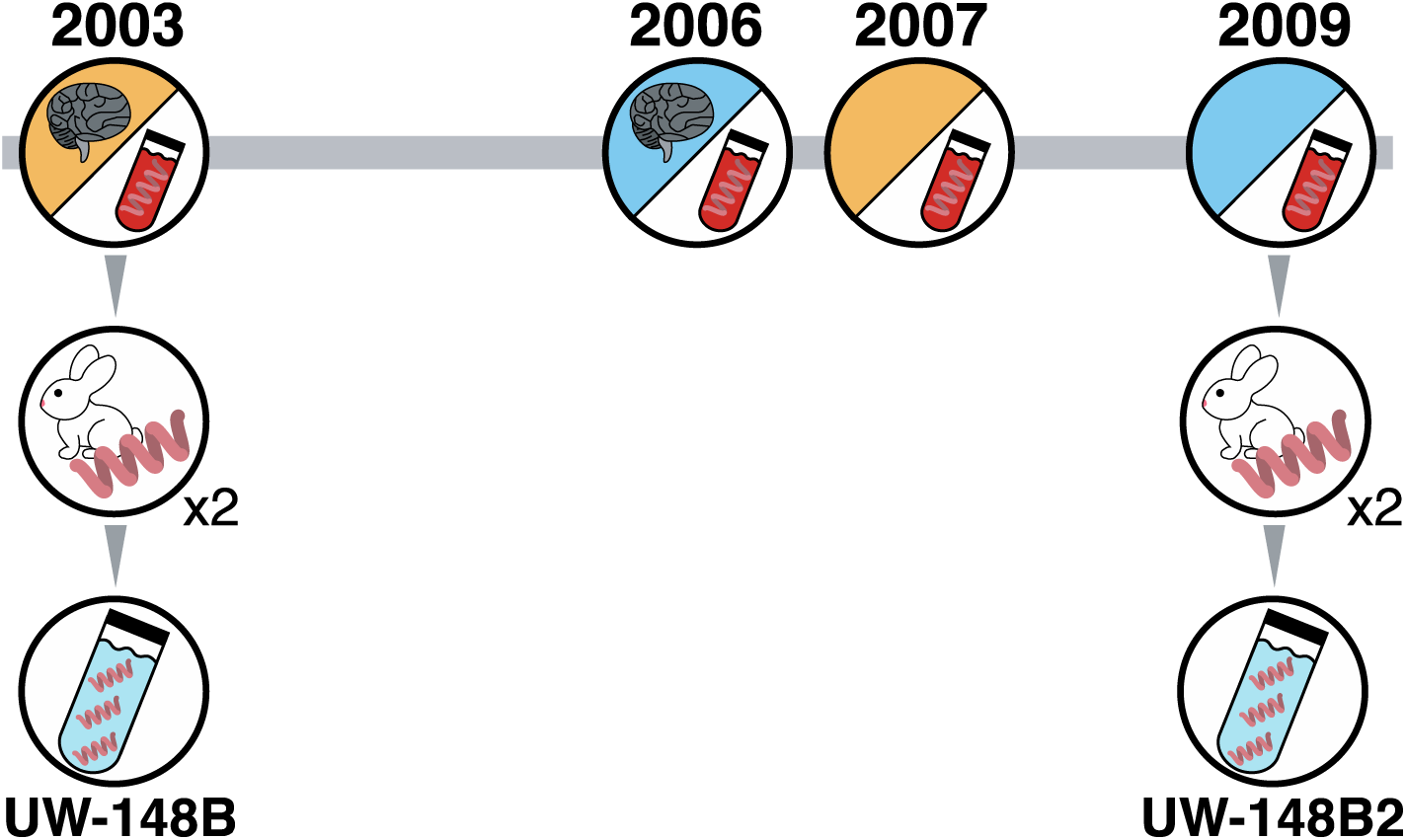
Isolation and propagation of UW-148B and UW-148B2. The two isolates in this study derived from the same patient who was exposed to *T. pallidum* four times over a six-year period. The stage of infection is indicated by the shading of the circle with secondary syphilis in orange and early latent syphilis in blue. Neurosyphilis is also indicated.

CSF-VDRL was nonreactive 3 months after treatment and serum RPR reverted to nonreactive a year after treatment. He presented with early latent syphilis in 2006, again had a reactive CSF-VDRL and was treated for neurosyphilis. CSF-VDRL reverted to nonreactive 3 months after treatment and serum RPR dropped from 1:2048 to 1:4 six months after diagnosis. He was treated for secondary syphilis without neurosyphilis in 2007 with serum RPR 1:256, and for early latent syphilis without neurosyphilis in 2009 with serum RPR 1:1024. Quantitative treponemal load testing of this episode yielded a median of 191 (range: 134 – 858) copies of *T. pallidum* DNA per mL of blood, of which 1mL was injected into a rabbit and passed a second time to derive the strain UW-148B2.

Treponemes harvested from the second rabbit for each strain were quantified by qPCR. The bioburden measured 3.64 × 10^6^ copies of *T. pallidum* DNA per mL of testicular harvest for UW-148B and 1.48 × 10^7^ copies per mL of harvest for UW-148B2. Using the enhanced molecular typing scheme with *arp, tprE, tprG*, and *tpr*J, and *tp0548*, the UW-148B yielded a strain type of 14d/f and the UW-148B2 yielded a 14d/g type [31].

### Assembly of complete genomes of UW-148B and UW-148B2

A total of 1.27 × 10^5^ and 5.17 × 10^5^ copies of *T. pallidum* DNA were used as input for capture sequencing for UW-148B and UW-148B2, and *de novo* assembly of the capture sequencing reads yielded 5 and 8 contigs that were greater than 15 kb. These contigs were aligned to the *T. pallidum* SS14 reference sequence (CP000805.1) to determine their positions within the genome. PCR and amplicon sequencing of the gapped regions were performed to close the genomes of the two isolates.

The *acidic repeat protein* gene (*tp0433*) in the UW-148B and UW-148B2 assemblies initially measured 600 bp and 540 bp, shorter than that of *T. pallidum* SS14. To determine if the difference in size of *tp0433* was the result of misassembly through these repetitive regions, we separately amplified and Sanger sequenced *tp0433*. Sequencing of *tp0433* revealed the genes in UW-148B and UW-148B2 were identical in size to that of *T. pallidum* SS14.

The complete genomes for UW-148B (CP045005.1) and UW-148B2 (CP045004.1) were 1,139,562 bp and 1,139,560 bp.

### Comparative genomics reveals parsimonious changes in outer membrane proteins between initial and reinfecting strains

A comparison of the genomes of UW-148B and UW-148B2 yielded 20 variants, excluding those found in the highly variable *tprK* gene (Fig 2; Table 1). Of these, 10 were single nucleotide polymorphisms (SNPs). Eight of these SNPs resulted in coding changes, which occurred in *tp0134, tp0151, tp0325, tp0326, tp515, tp0548 and tp0691*, between the two strains, while the remaining two, located in *tp0319* and *tp0404*, were synonymous mutations.

**Table 1.** Genomic mutations distinguish UW-148B and UW-148B2.

| Nucleotide Position (CP045005.1) | Gene | Nucleotide Change | Amino Acid Change | Protein Effect | Present in both shotgun and capture sequencing? |
| --- | --- | --- | --- | --- | --- |
| 12485 | *tp0012* | 110delG | Gly37Valfs*17 | Restored ORF | Yes |
| 34084 | - | insG | - | Intergenic | Yes |
| 49369 | *tp0040* | 2419delG | Val807* | Truncation | Yes |
| 140956 | - | insC | - | Intergenic | Yes |
| 156523 | *tp0134* | 233G>A | Thr78Ile | Substitution | Yes |
| 158621 | *tp0136* | 251delG | Met1ext-122 | Restored ORF | Yes |
| 174174 | *tp0151* | 778C>T | Val260Ile | Substitution | Yes |
| 208630 | - | insG | - | Intergenic | Yes |
| 336944 | *tp0319* | 930T>C | - | None | Yes |
| 342698 | *tp0325* | 1618G>A | Ala540Thr | Substitution | Yes |
| 346246 | *tp0326* | 730A>G | Thr244Ala | Substitution | Yes |
| 409148 | - | delG | - | Intergenic | No |
| 430384 | *tp0404* | 198T>C | - | None | Yes |
| 510331 | *tp0479* | delC | Met1ext-39 | Restored ORF | Yes |
| 531771 | *tp0496* | insT | Met1ext-15 | Restored ORF | Yes |
| 555552 | *tp0515* | 1366C>T | Arg458Cys | Substitution | Yes |
| 592692 | *tp0548* | 28G>A | Gly10Arg | Substitution | Yes |
| 592696 | *tp0548* | 32G>A | Gly11Glu | Substitution | Yes |
| 759489 | *tp0691* | 83T>C | Lys28Arg | Substitution | Yes |
| 937290 | *tp0859* | 641delG | Gly214Valfs*16 | Re-split fused ORFs | Yes |

**Fig 2.**
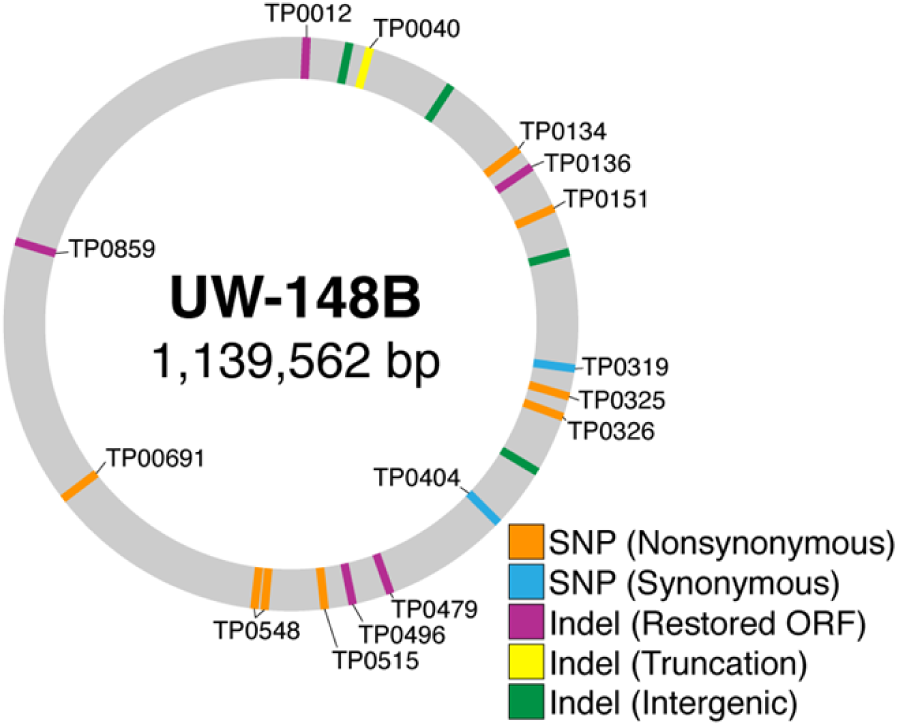
Variants outside of *tprK* distinguish initial and reinfecting isolates. Genomic mutations and their impact on open reads frames (ORFs) in the reinfecting strain (UW-148B2) relative to the initial strain (UW-148B) are denoted. Eight non-synonymous SNPs and two synonymous SNPs were recovered along with ten indels.

In addition to SNPs, there were 10 insertions or deletions in homopolymeric repeats. Six of these changes occurred within open reading frames (ORFs) and the remaining 4 were in intergenic regions. The homopolymeric mutation in *tp0040* resulted in a truncation in UW-148B2, while the other 4 mutations restored ORFs, including *tp0012, tp0136, tp0479 and tp049*6, in UW-148B2 that were disrupted in UW-148B. Notably, the ORFs of *tp0859* and *tp0860* were fused in UW-148B. A homopolymeric deletion in UW-148B2 split these two genes.

### The reinfecting strain contains less diversity in TprK compared to initial strain

To understand diversity of the highly variable *tprK* gene, we sequenced the 1.6kb gene using both short- and long-read approaches. When generating the full-length *tprK* amplicons for both short- and long-read sequencing, a total of 7.28 × 10^3^ and 2.96 × 10^4^ copies of *T. pallidum* DNA for UW-148B and UW-148B2 were used as input template for PCR and sequenced to an average depth of 13,366x and 14,404x. We first examined diversity within each of the variable regions using short-read sequencing of the *tprK* amplicon. To account for potential sequencing error, we required that each unique amino acid sequence within a given variable region be supported by at least 5 reads (S2 Table). We identified 4 SNPs resulting in 2 coding changes compared to the *tprK* gene of *T. pallidum* SS14 in the 13 bases 5’ to the previously defined V3 variable region, suggesting the V3 region extends outside of the current SS14 *tprK* annotation (CP000805.1). For subsequent analyses, we extended our definition of the V3 variable region to include this region.

After quality filtering, the number of unique amino acid sequences in the 7 from which variable regions ranged from 11 – 140 sequences for UW-148B and from 8 – 107 sequences for UW-148B2 (Fig 3). For both isolates, V6 displayed the most sequence heterogeneity, while V1 was the least heterogeneous. We also observed considerable variation in the lengths of V3, V6 and V7 sequences within each isolate. The difference in size between the longest and shortest variants of V3, V6 and V7 were 15, 16, and 17 amino acids for UW-148B and 7, 14, and 18 amino acids for UW-148B2. Of note, just one variant from all 7 variable regions in UW-148B produced an internal stop codon in the Illumina sequencing data using a support threshold of 5 reads. This variant was located in V4 and supported by only 5 reads. UW-148B2 did not contain any variants with stop codons.

**Fig 3.**
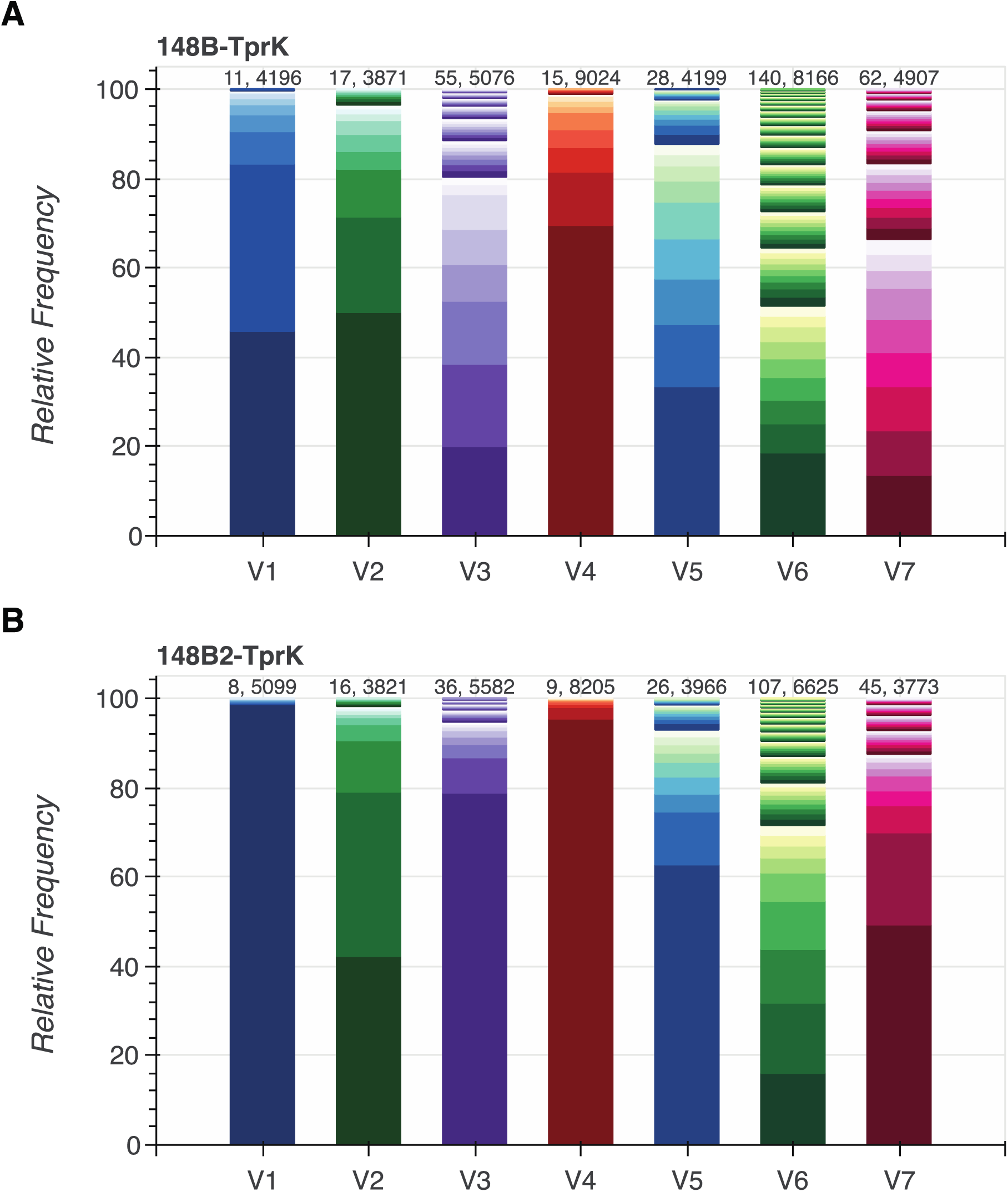
Relative frequencies of the different amino acid sequences identified in each of the 7 variable regions (V1 – V7) of TprK. Relative frequencies of the amino acid sequences for each of the variable regions identified through short-read sequencing of TprK for A) the initial strain (UW-148B) and B) the reinfecting strain (UW-148B2). The number of sequences and total number of reads spanning a given variable region are displayed above each bar.

Next, we compared the heterogeneity of each of the variable regions between the initial and reinfecting strains. UW-148B contained more variants for every variable region compared to UW-148B2 (Fig 3). We then applied Pielou’s evenness measure to determine if the observed variants of each variable region were equitably distributed. Each of the variable regions of UW-148B was more evenly distributed than the corresponding variable region of UW-148B2 (Table 2). Next, we calculated the Shannon diversity index for each variable region of each strain. Reflecting the higher number of variants and more even distribution of variants, each of the variable regions of UW-148B was more diverse than those of UW-148B2 (Table 2).

**Table 2.**
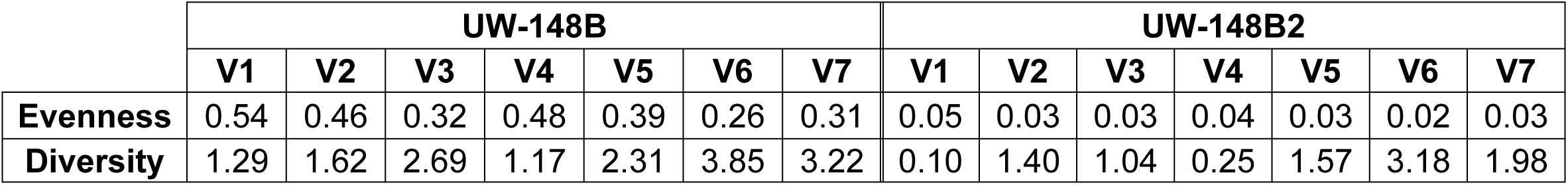
Pielou’s evenness and Shannon diversity measures calculated for each of the variable regions of the two strains.

|  | UW-148B |  |  |  |  |  |  | UW-148B2 |  |  |  |  |  |  |
| --- | --- | --- | --- | --- | --- | --- | --- | --- | --- | --- | --- | --- | --- | --- |
|  | V1 | V2 | V3 | V4 | V5 | V6 | V7 | V1 | V2 | V3 | V4 | V5 | V6 | V7 |
| **Evenness** | 0.54 | 0.46 | 0.32 | 0.48 | 0.39 | 0.26 | 0.31 | 0.05 | 0.03 | 0.03 | 0.04 | 0.03 | 0.02 | 0.03 |
| **Diversity** | 1.29 | 1.62 | 2.69 | 1.17 | 2.31 | 3.85 | 3.22 | 0.10 | 1.40 | 1.04 | 0.25 | 1.57 | 3.18 | 1.98 |

### UW-148B and UW-148B2 share variants in their variable regions at different frequencies

We compared the amino acid sequences in each of the variable regions of each strain to assess whether the two strains shared sequences. In total, the two strains contained 41 identical individual variable sequences across all of the variable regions. Notably, the two strains did not share any sequences in V3. Next, we examined the difference in relative frequencies of the shared variants between the two strains. For UW-148B, the highest frequency variants of V1, V4, V5 and V7 were present in UW-148B2, albeit at lower frequencies (Fig 4). The highest frequency variants of V2 and V6 for UW-148B were not present in UW-148B2. Similarly, the highest frequency variants of V1, V2 and V4 from UW-148B2 were present at lower frequencies in UW-148B. The highest frequency V5, V6, and V7 variants of UW-148B2 were not present in UW-148B.

**Fig 4.**
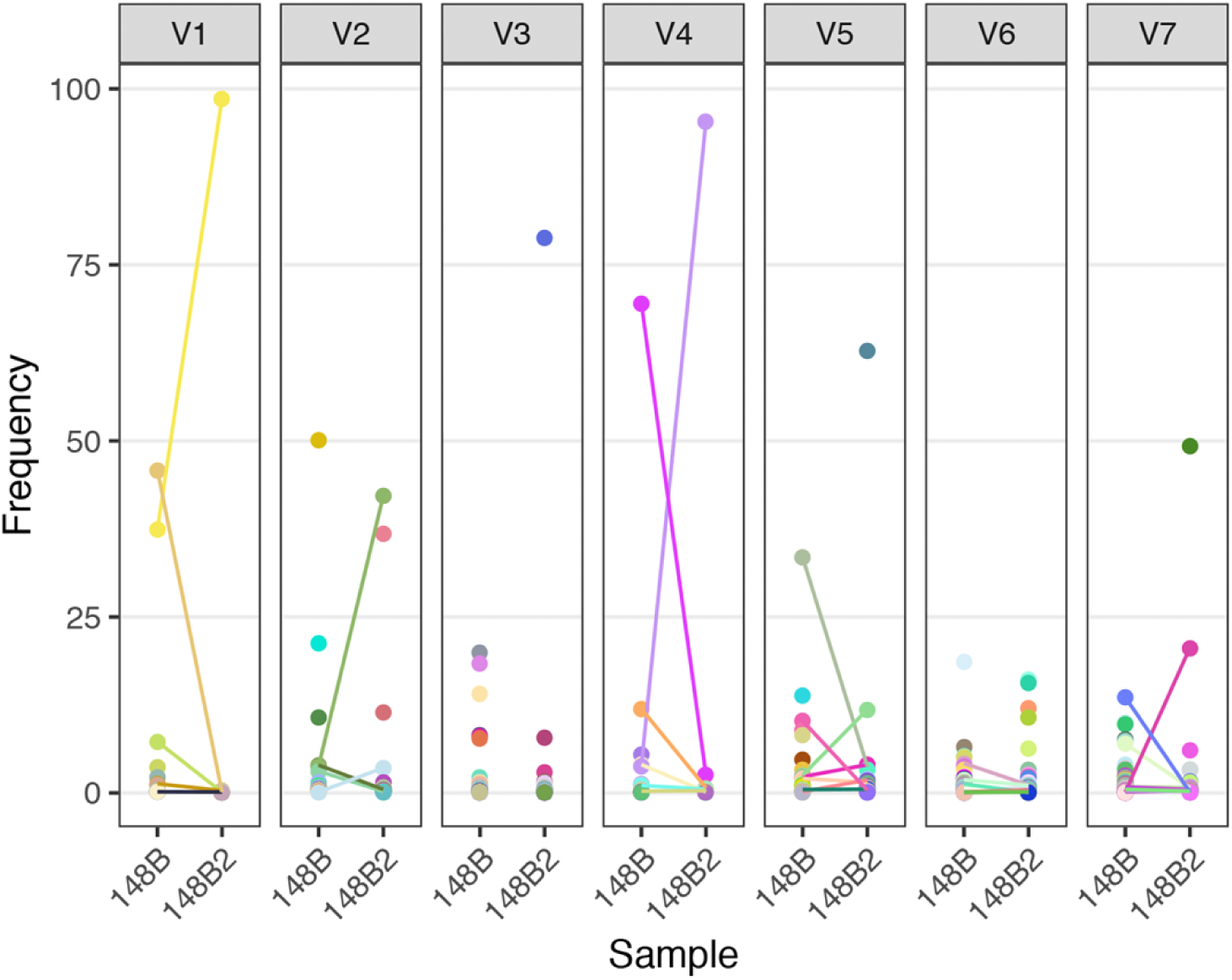
Comparison of the sequences from each of the variable regions identified in the initial and reinfecting *T. pallidum* strains. Illumina-based frequencies of each unique amino acid sequence for each variable region within TprK are shown. Sequences are matched by color within the same variable region and dots connected by lines denote sequences present in both UW-148B and UW-148B2. Unconnected dots are present in only the depicted sample.

### Long-read sequencing reveals UW-148B and UW-148B2 do not share identical copies of TprK

As long-read sequencing often produces more errors than short-read sequencing, we used a read clustering-based denoising approach [25] to identify probable sequences of the *tprK* amplicons for each isolate. We filtered Q20 PacBio CCS reads based on size (1,400 - 1,800 bp) prior to applying denoising and then required each cluster to contain a minimum of 5 reads. We compared the sequences of the variable regions obtained from these denoised sequences to those obtained from short-read sequencing of the *tprK* amplicon to determine the accuracy of long-read sequencing and our quality filtering. This comparison revealed that short-read and denoised long-read sequencing results were consistent (R^2^: 0.990 and 0.991 for UW-148B and UW-148B2; Fig 5). For UW-148B, all the sequences identified from short-read sequencing at a frequency above 0.062% were also found in the long-read data. For UW-148B2, all the sequences from short-read sequencing at a frequency above 1.09% were identified in the long-read data.

**Fig 5.**
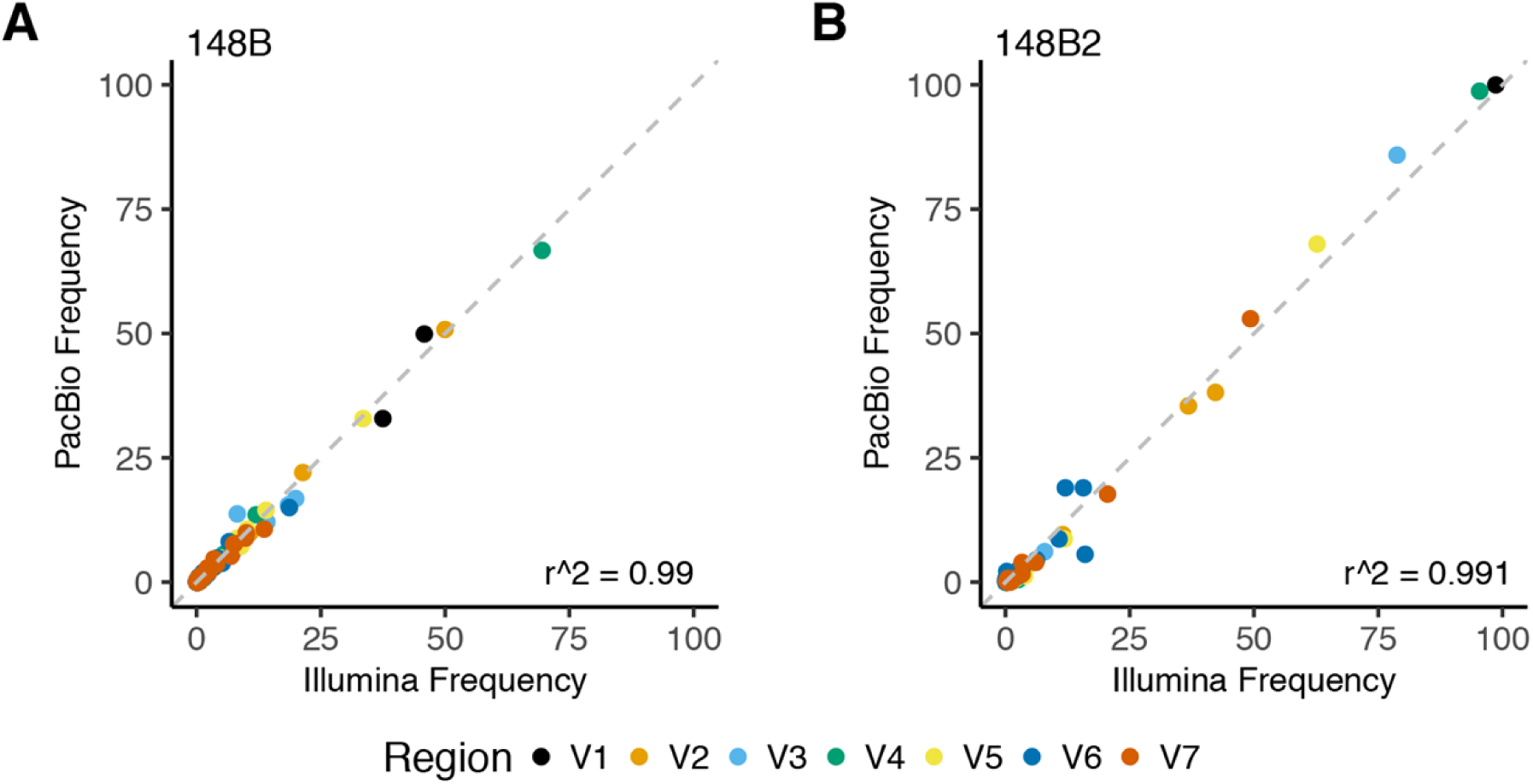
Comparison of short-read and long-read sequencing of TprK. Comparison of the determination of identical amino acid sequences in each of the variable regions of TprK through short-read (Illumina) and long-read (PacBio CCS) sequencing approaches for A) the initial strain, UW-148B and B) the reinfecting strain, UW-148B2.

For both isolates, long-read sequencing generated sequences in each of the variable regions that were not found in the short-read data. These sequences contained internal stop codons or resulted in incorrect translation of the constant regions of TprK and were present at low frequencies. Thus, we excluded full-length sequences present at a frequency less than 0.2% from the long-read data for subsequent analyses (S3 Table).

Next, we examined the relationship between the TprK sequences from the initial and reinfecting strain. UW-148B contained more full-length TprK sequences at a frequency greater than 0.2% compared to UW-148B2 (127 sequences versus 114 sequences). We then constructed a phylogenetic tree with the TprK sequences from both strains. The TprK sequences of each isolate segregated based on the strain from which they were derived from (Fig 6). Notably, the two strains did not share any identical full-length TprK sequences.

**Fig 6.**
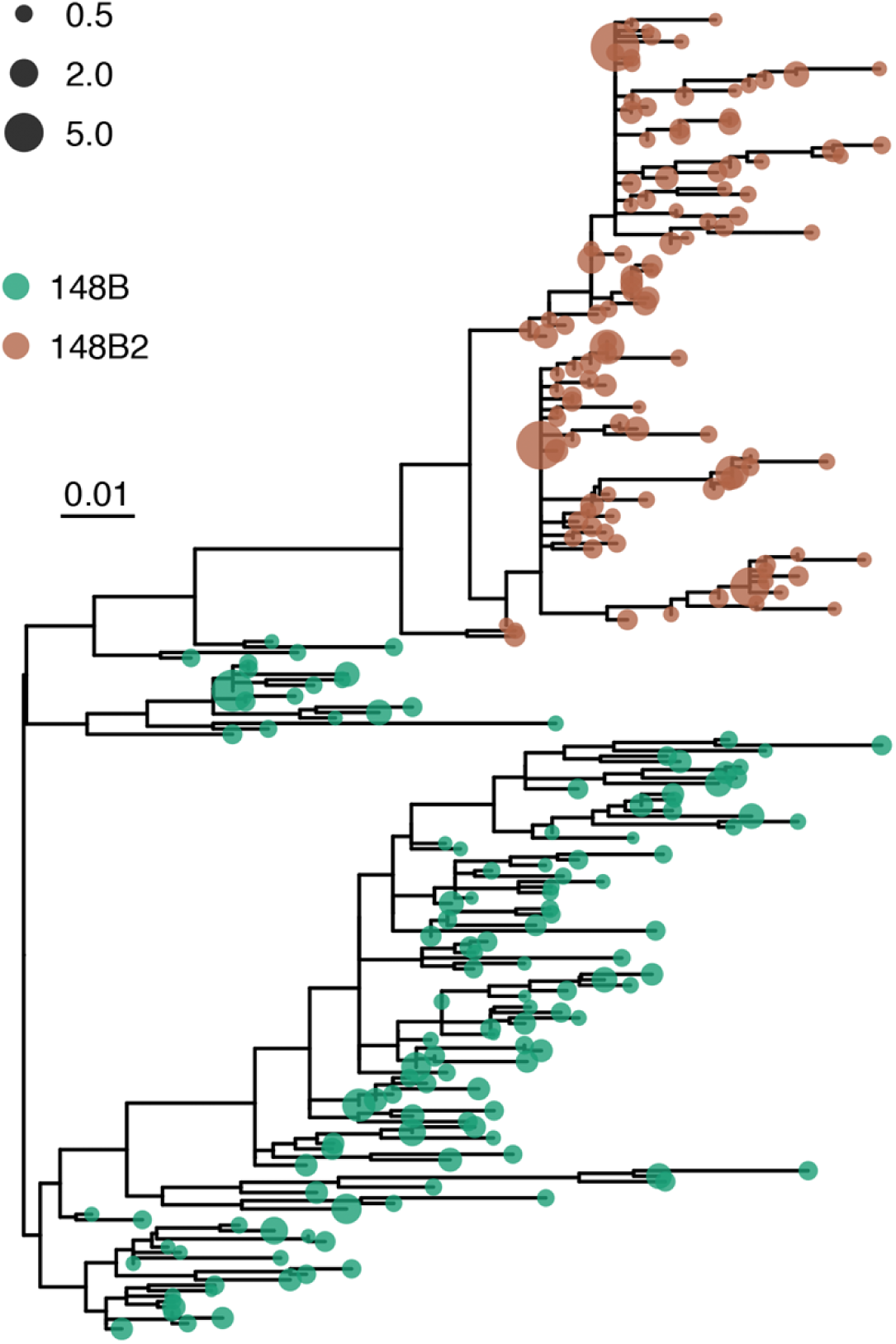
Phylogenetic tree constructed with the full-length TprK sequences of the initial and reinfecting strains. The color of each of node depicts the strain from which the full-length TprK sequence originated. A green node depicts those sequences from the initial strain, UW-148B, while a bronze node depicts those from the reinfecting strain, UW-148B2. The size of each node indicates the relative percentage of the full-length TprK sequence in the strain from which the sequence originated.

## Discussion

Here, we present a comparison of complete genomes with deep, full-length *tprK* profiling, of two *T. pallidum* subsp. *pallidum* isolates derived 6 years apart from a single patient. We demonstrated that limited genomic changes distinguish the two isolates. Using both short- and long-read sequencing to investigate the changes occurring in the highly variable *tprK* gene, we show that regions of TprK are more diverse in the initial isolate as compared to the reinfection isolate. Long-read sequencing yielded the first complete sequences of TprK at a depth that demonstrates the high level of intra-isolate heterogeneity of this protein. We further determined that the two isolates, despite sharing certain variants within a given variable region, did not share a single identical full-length sequence of TprK.

The global *T. pallidum* subspecies *pallidum* population shows a remarkable lack of genetic variation outside of *tprK* [21]. The initial and reinfecting isolates analyzed in this study reflect this lack of genomic diversity that exists in the global *T. pallidum* population when *tprK* sequences are not considered. The two strains were only distinguished by 20 genomic changes outside of *tprK*. Of those, 14 resulted in coding changes in 13 genes. Notably, all of the coding changes found in the reinfecting strain were identified in previously sequenced strains [13,21,32] and thus, likely reflect the diversity in the circulating *T. pallidum* population rather than selection from the host’s immune response.

Intriguingly, 9 of the 14 genes with coding differences between the two strains have been demonstrated or predicted to encode for outer membrane or periplasmic proteins [33–41]. These include TP0040, TP0134, TP0136, TP0151, TP0325, TP0326, TP0515, TP0548 and TP0859. Two of these proteins, TP0136 and TP0326, are known targets of the host immune response in the rabbit [36,39,40,42,43].

TP0136 encodes an outer membrane protein responsible for binding to the host’s extracellular matrix [40,42]. The N-terminal portion of the protein, in particular, plays a significant role in binding to plasma fibronectin [40]. Immune responses targeting TP0136 can delay lesion ulceration [42] and inhibit bacterial binding to fibronectin [40]. Interestingly, the initial isolate in our study had a homopolymeric insertion, which shifted the start codon of TP0136 downstream and likely disrupted the N-terminal of the resulting protein. Further study is required to understand how this mutation impacts protein expression and stability as well as the immune response to TP0136.

TP0326 encodes a protein belonging to the β-barrel assembly machinery A (BamA) protein family [39]. In both humans and rabbits, this protein elicits a strong immune response [36,39,43]. Antibodies target the surface-exposed loops of the β-barrel, as well as the polypeptide transport-associated (POTRA) domains of TP0326 [39,44]. However, a stronger response is mounted against the POTRA domains [44]. The coding change seen between the two isolates here (T260A) mapped to the POTRA3 domain. While the impact of amino acid substitutions in the TP0326 β-barrel on immunoreactivity has been studied [44,45], further work is required to understand the impact of the amino acid substitution observed in our isolates.

Previous studies using next-generation sequencing to investigate the outer membrane protein TprK have shown that a significantly greater intra-isolate diversity exists within the 7 variable regions of *tprK* compared to the rest of the genome [20,46]. Our two isolates also contained a considerable amount of diversity in each variable region. Also consistent with previous findings [20,46], V6 was the most diverse region, while V1 was the least diverse region in both strains. Unlike previously sequenced isolates [20], our isolates had more variants present at frequencies between 20-80%. This difference may be attributable to our isolates’ two passages in rabbits, resulting in high treponemal loads along with immune pressure in the rabbit, or may reflect variations in TprK during different stages of syphilis infection.

In rabbit models, antibodies are developed against V2, V4, V5, V6 and V7 of TprK [19]. These antibodies are specific to a single variant of a variable region [19,47]. While our strains shared certain variants of each variable region, these variants were present in different frequencies in the two strains. Additionally, the reinfecting strain showed less diversity in each of the 7 variable regions compared to the initial strain. These differences may reflect the development of host immunity (either in the human or rabbit) against variants over time, constraining the potential diversity of TprK in the reinfecting strain.

We present the first full TprK sequences at a depth that highlights the considerable intra-isolate heterogeneity of this protein using long-read sequencing. Our results indicate long-read sequencing is remarkably consistent with short-read sequencing for detecting TprK variants, consistent with the increasing accuracy of long-read sequencing [48]. However, as long-read sequencing is still less accurate than short-read sequencing [25,49], we could not confidently call TprK variants present at lower frequencies (< 0.2%). Increasing sequencing depth may help resolve TprK variants present at frequencies less than 0.2%. Our approach to discerning the full sequence of the TprK variants present in a given strain may also prove useful for other organisms containing highly variable genes, including *Borrelia burgdorferi* [50] and *Neisseria gonorrhoeae* [51]. The ability to profile full-length TprK sequences will be critical for developing TprK typing schemes, nomenclature, and databases, as well as serological studies with programmable peptide microarrays and phage display libraries to fully characterize the anti-TprK immune response.

Our study was limited by the passage in rabbits used to amplify the *T. pallidum* strains. Until recently, an *in vitro* culture system did not exist for *T. pallidum* [52] and passage through rabbits was helpful for obtaining sufficient treponemal genetic material for whole-genome sequencing. As a result, the genetic changes we observed between the two strains may reflect adaptations to the rabbit or, in the case of TprK, gene conversion that occurred in the rabbits, and thus may not be fully reflective of *T. pallidum* sequence in humans. Passage in rabbits could result in increased measurements of TprK diversity by allowing for additional cycles of gene conversion and/or purifying immune selection.

We were further limited by our sample size as isolating reinfecting strains can prove difficult [53]. Further work examining the genomic changes associated with reinfection may benefit from the *in vitro* culture system or, in the case of patient samples with higher treponemal levels, capture sequencing. In addition, our analysis of TprK diversity within the two strains was limited by introduced population bottlenecks while collecting genetic material from each strain. Our estimates of diversity within the reinfecting strain, in particular, may be limited by the nearly 13-fold lower inoculation of treponemes into the first rabbit. Future studies will be required to understand the rate of full-length TprK evolution in both humans and the rabbit model as well as how inoculum loads and copy number affect diversity measurements. Despite the limitations of the rabbit model, the inability to find the same full-length TprK sequence across more than 100 TprKs in each of the two strains is illustrative of the impressive ability of this protein to present antigenic diversity and consistent with a model of ongoing selection and winnowing of TprK diversity over time in reinfection.

As rates of syphilis continue to steadily increase, a vaccine against *T. pallidum* is a public health necessity [4]. Examining the genomic changes in *T. pallidum* that permit reinfection, as well as profiling the diversity existing within the putatively immunodominant outer membrane protein TprK, is critical for guiding the development of an effective vaccine. Our results indicate conservative mutations in outer membrane or periplasmic proteins may also represent a genetic basis through which *T. pallidum* escapes cross-protective immunity; however, immunological confirmation in future studies is required. Our results also highlight the critical need to target conserved regions of outer membrane proteins for vaccine antigens given the highly mutable immunoevasion mechanism of *tprK*.

## Supporting Information Legends

**Supplemental Table 1.** PCR primers used in this study for gap filling and deep tprK profiling.

**Supplemental Table 2.** Variable region sequences identified in tprK of UW-148B and UW-148B2 from either Illumina or denoised Pacbio sequencing, filtered to require 5 reads of support.

**Supplemental Table 3**. Full-length TprK sequences identified in UW-148B and UW-148B2 with greater than 5 reads of support after denoising and frequency of >=0.2%.

